# A psychophysical characterization of tactile and temperature perception on the human torso

**DOI:** 10.64898/2026.09.30.755664

**Authors:** Sophia Faresse, Roger Holmes Watkins, Rochelle Ackerley

## Abstract

Numerous studies have been conducted to investigate touch on the hand and forearm, but less is known about the rest of the body, particularly the torso and breasts, which play an important role in affective touch. We conducted psychophysical tests on three skin sites on the torso: upper breast, nipple-areola complex (NAC), and under the breast, as well as on two control sites: palm and forearm. We studied touch pleasantness and intensity by applying gentle caress-like strokes using a soft brush, and measured temperature and tactile detection thresholds. The results show an inverted U-shaped relationship for pleasantness over increasing stroking velocity for all skin sites, although we found some differences between sites, particularly at 30 cm.s^-1^, where participants found stroking on the NAC to be more intense and women additionally found it to be less pleasant. For temperature detection, we found that the NAC was more sensitive in men. For tactile detection, we mainly observed that the areola was far less sensitive than the other areas. Together, these results provide insights into tactile perception in the torso on a fundamental level, which can be applied to understanding how to regain sensations after mastectomy.

## Introduction

The sense of touch plays a crucial role throughout life: it allows us to distinguish objects and navigate our environment, but it is also a mode of non-verbal communication, crucial in human-to-human interactions (Gallace and Spence 2010). Although the hand is primary in exploratory touch, when it comes to emotional and intimate exchanges, the torso and breasts play a major role. One of the common forms of tactile communication is hugging, which has positive effects on stress regulation (Light et al. 2005). Further, touch is particularly involved in intimate relationships and sexuality, with breasts being erogenous zones (Turnbull et al. 2014) of high capabilities for sexual arousal (Nummenmaa et al. 2016). However, even though the torso and breasts are important in human non-verbal communication, we know relatively little about their tactile and thermal sensitivity, and the underlying physiology, as compared to the hand.

Touch is mediated through low-threshold mechanoreceptors (LTMRs), located at the end of afferent nerve fibers in the skin, and decades of research have identified several distinct classes that fire optimally to specific types of touch. Fast-conducting Aβ-LTMR fibers have been found to be essential for the awareness of touch and tactile discrimination, playing a key role in the determination of what touched us, where, and for how long (McGlone et al. 2014). A specific type of low-threshold C-fiber, named C-LTMR afferents, or C-tactile (CT) afferents in humans, also exist and are postulated to reinforce the positive affective aspects of touch (McGlone et al. 2014), as their activity correlates with pleasantness ratings, where gentle stroking at 1-10 cm.s^-1^ around skin temperature is optimal (Löken et al. 2009; Ackerley, Wasling, et al. 2014). C-LTMRs form the basis of the affective touch hypothesis (Vallbo et al. 2009; Olausson et al. 2010; Morrison et al. 2010), which also led to the social touch hypothesis (Morrison et al. 2010), theorizing that they specifically reinforce human-to-human interactions. Human in vivo microneurography studies have found that C-LTMRs are dense in the hairy skin of the forearm, but rare in the glabrous skin of the hand (Watkins et al. 2021). Further, they have been found to be present on the face (Nordin 1990; Johansson et al. 1988) and relatively sparse on the leg (Löken et al. 2022). Although comparative work suggests that the equivalent C-LTMRs in animals are denser proximally (Li et al. 2011), there is insofar no direct evidence available to support or contradict this in humans.

Studies have attempted to link the perception of stroking touch (Ackerley, Carlsson, et al. 2014; Cazzato et al. 2021; Jönsson et al. 2015), and even vicarious (observed) touch (Morrison et al. 2011; Schirmer and McGlone 2019; Ali et al. 2023; Bellard et al. 2022; Walker et al. 2017), to the density of C-LTMRs in humans; however, these results cannot provide direct evidence of C-LTMR density and can only imply it through the shape of the pleasantness curve over different stroking velocities. The pleasantness curve can be heavily influenced from other factors (e.g. temperature, odors) (Ackerley, Wasling, et al. 2014; Croy et al. 2014), where there can be a divergence between pleasantness perception and C-LTMR activity (Ackerley, Wasling, et al. 2014). Further, studies comparing the pleasantness of stroking on the C-LTMR-dense forearm and glabrous hand skin, where C-LTMRs are rarely found, find few differences (Ackerley, Carlsson, et al. 2014; Cazzato et al. 2021; Cruciani et al. 2021). This raises questions as to the strength of the coupling between tactile pleasantness and C-LTMR activity, where pleasantness can nevertheless be high and show an inverted-U shaped relationship (Ackerley, Carlsson, et al. 2014; Cazzato et al. 2021; Jönsson et al. 2015; Cruciani et al. 2021), even with relatively lower C-LTMR density (Watkins et al. 2021; Löken et al. 2022), thus relying on other inputs (e.g. Aβ-LTMRs).

Psychophysical tactile pleasantness studies, stroking the skin over different velocities, have shown that areas on the face and limbs show relatively similar inverted-U shaped pleasantness profiles for all the skin sites, although there is a lot of variability within- and between-studies (Cruciani et al. 2021). To our knowledge, little work has investigated tactile pleasantness on the torso, with one recent study finding that stroking touch on the lower abdomen was perceived as less pleasant than on the palm, forearm, and face (Cazzato et al. 2021), which is somewhat contrary to the proximal C-LTMR density proposition. Studies using vicarious touch have shown that watching someone being stroked on the back is very pleasant (Walker et al. 2017; Haggarty et al. 2021), although this increase was not found in actual physical stroking (Bellard et al. 2022). As well as being implicated in pleasant touch, C-LTMRs have also been postulated to underlie erotic touch (Jönsson et al. 2015; Bendas et al. 2017), although these were not conducted on the torso.

Akin to pleasant touch research, few studies have investigated other tactile sensations on the torso, but due to its important role in body-self and social interactions, investigations of tactile perception on the torso, especially on the breast, have gained recent interest. Long et al. (2025) studied the spatial acuity of the breast and compared it to that of the hand and back. They found that the spatial acuity of the breast, especially the nipple-areola complex (NAC) was low, even lower than that of the back, which is one of the lowest areas on the body (Weinstein 1968). Further, they found that spatial acuity decreased with breast size and that the nipple was perceived as a sensory unit, meaning that the localization of sensation on the breast was biased toward the nipple (Long et al. 2025). Tactile research on the breast is particularly important, due to its role in affective and erogenous touch and recent work has postulated the potential for a Bionic Breast, a sensory neuroprosthetic after mastectomy (Lindau and Bensmaia 2020). The breast is a unique skin area, containing hairy, glabrous, and mucocutaneous skin; thus, its innervation by sensory afferents is likely complex. Understanding sensory perception on the breast is therefore important for advancing general tactile research, but also in its application to neuroprosthetics.

Recently, researchers studied the normative values of nipple areola complex (NAC) sensation, measuring tactile detection thresholds and two-point discrimination thresholds on five sites: the nipple and the areola divided into four quadrants (Kasielska-Trojan et al. 2021). They found that the nipple was the most sensitive part, and that tactile detection thresholds correlated with age, body mass index, history of births, breast size, breast ptosis, breastfeeding history, and areola diameter. More recently, a study was conducted on women diagnosed with breast cancer to assess breast sensibility in non-operated and operated breasts, measuring tactile detection thresholds and thermal detection thresholds on the NAC and the four breast quadrants. They found the NAC to be less sensitive than the breast quadrants, and impaired sensation after breast surgery, especially after mastectomy, both for tactile detection and thermal detection (Bubberman et al. 2025). Thus, breast surgery clearly affects sensation and there is the potential to ameliorate this.

One other aspect of innocuous somatosensation is temperature sensing. Touch always has a thermal element – even if the contact is at neutral skin temperature – and touch without temperature would be like vision without color. Temperature greatly modifies affective touch (Ackerley, Wasling, et al. 2014; Cabanac 1971; Marks and Gonzalez 1974) and we all know from personal experience that cold touch on the abdomen is unpleasant. In general, temperature changes are more readily detected on the upper half of the body, particularly the face, and cooling is overall more easily perceived than heating (Stevens and Choo 1998). It has been shown that the perception of cooling varies significantly across the torso in men, with the lower and lateral abdomen being more sensitive to cold (Filingeri et al. 2014). More recently, researchers have focused specifically on temperature perception on women’s breasts and have measured the perception of static thermal stimuli, finding that the perception of warm stimuli, but not cool, varied significantly across the breast (Valenza et al. 2023).

Overall, it is evident that tactile and thermal sensations on the breast warrant more investigation, especially comparisons between other skin sites where the sensory innervation is highly different. Presently, we aimed to further characterize touch perception on the breast and torso, in comparison to control sites of the glabrous palm and hairy forearm, particularly focusing on pleasant touch perception. To this end, we conducted a psychophysical study divided into two parts: (1) In the first part, we used a classic stroking affective touch protocol, as delivered by a robot, to investigate the relationship between pleasantness perception and stroking velocity on five skin sites, three on the torso: upper breast, NAC, and non-breast torso, and two control skin sites: the forearm and the palm. In addition, we investigated thermal sensitivity by measuring thermal detection thresholds to warm and cool. In the second part, we conducted a complementary affective touch protocol to compare affective touch perception of manual brush stroking stimulation delivered by a trained experimenter across the same skin sites except for the NAC. In addition, we investigated tactile sensitivity by measuring tactile detection thresholds to force indentation on the four skin sites, plus on the nipple and on the areola separately. We hypothesized that we would find a negative quadratic relation between pleasantness rating and stroking velocity and a linear one for tactile intensity, as previously shown for other skin sites (Löken et al. 2009; Ackerley, Carlsson, et al. 2014; Jönsson et al. 2015; Sehlstedt et al. 2016; Triscoli et al. 2013) and we predicted that due to the complexity of the NAC, this would show particular differences in perception.

## Methods

### Ethics

Our study was approved by the CPP Paris-Ile de France X ethics committee under the number 2023-A01990-45. All procedures were following the 1964 Helsinki Declaration and its later amendments apart from pre-registration in a database. Participants were recruited via mailing lists, flyers, and word of mouth. Exclusion criteria included current being pregnant or breastfeeding, having had surgery on the breasts, and having a disorder or condition that may alter tactile sensitivity (declarative, e.g. diabetic neuropathy, dermatological condition, psychiatric or neurological disorders). Informed consent was obtained from all participants included in this study. They were paid €10 per hour. Before the experiment, time was spent explaining the experiment, showing the device, and answering questions.

### General data collected over both experiments

Information collected for population description included age, sex, breast size (in cm, measured as the difference between the torso circumference on the breast (nipple level) and below the breast (inframammary fold in women, and subpectoral in men), areola size (in cm^2^, calculated according to the area of an oval based on the measurement of the rostrocaudal and mediolateral diameter of the areola), current use of hormonal contraception (Table 1). Breastfeeding history was also obtained, but only three women had previously breastfed, therefore this was not considered in the analyses. At the end of both experiments, participants filled the short version of the Touch Experiences and Attitudes Questionnaire (TEAQ-s) (Friedrich et al. 2025). The TEAQ-s is a 16 items self-reported questionnaire that assesses experiences and attitudes of positive touch recently and across the lifespan, with higher scores corresponding to more positive attitudes and experiences of touch. It comprises 4 dimensions (4 items each): attitude to friendly touch (AFT), current intimate touch (CIT), childhood touch (CHT) and attitude to intimate touch (AIT). The mean TEAQ-s scores for each experiment can be found in Table 1.

**Table 1:**
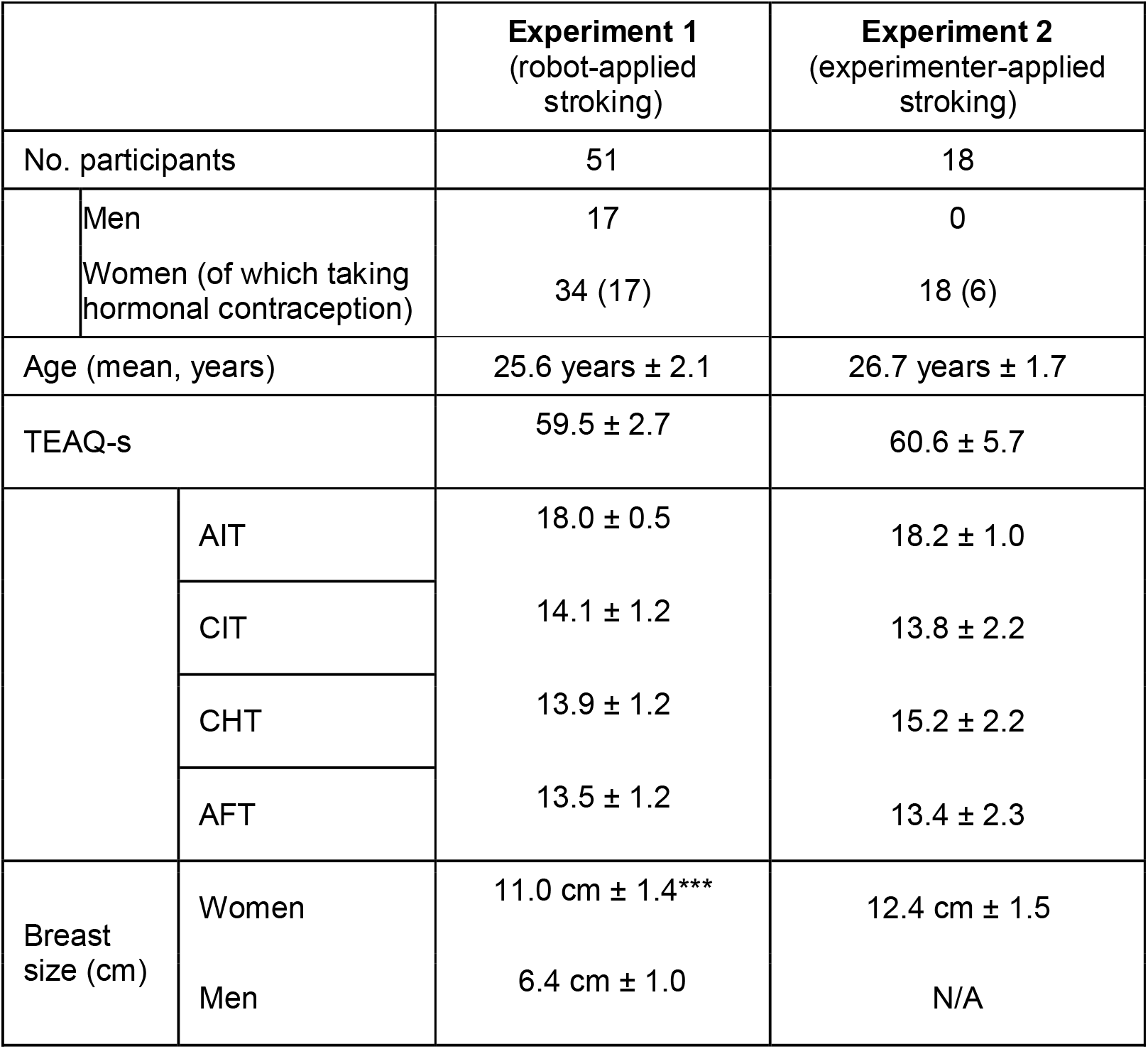
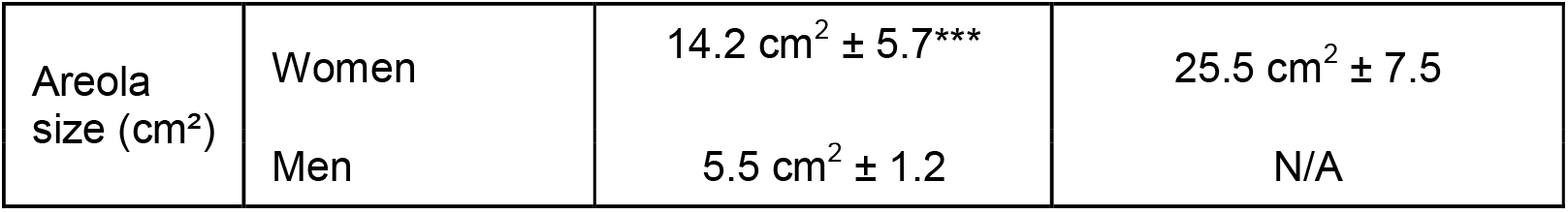
Characteristics of participants in Experiment 1 and Experiment 2. Number of participants are presented (n). Mean (M) scores and measures are presented +/-the 95% confidence interval. In Experiment 1, we tested differences between breast size and areola size between men and women using unpaired t-tests, finding that women had significantly larger breasts (t = 5.36, df = 49.0, p < 0.001) and significantly larger areolae (t = 4.49, df = 33.6, p < 0.001), as indicated by ^***^.

|  |  | **Experiment 1**<br>(robot-applied<br>stroking) | **Experiment 2**<br>(experimenter-applied<br>stroking) |
| --- | --- | --- | --- |
| No. participants |  | 51 | 18 |
|  | Men | 17 | 0 |
|  | Women (of which taking<br>hormonal contraception) | 34 (17) | 18 (6) |
| Age (mean, years) | | 25.6 years ± 2.1 | 26.7 years ± 1.7 |
| TEAQ-s | | 59.5 ± 2.7 | 60.6 ± 5.7 |
| | AIT | 18.0 ± 0.5 | 18.2 ± 1.0 |
| | CIT | 14.1 ± 1.2 | 13.8 ± 2.2 |
| | CHT | 13.9 ± 1.2 | 15.2 ± 2.2 |
| | AFT | 13.5 ± 1.2 | 13.4 ± 2.3 |
| Breast<br>size (cm) | Women | 11.0 cm ± 1.4*** | 12.4 cm ± 1.5 |
| | Men | 6.4 cm ± 1.0 | N/A |
| Areola size (cm<sup>2</sup>) | Women | 14.2 cm<sup>2</sup> ± 5.7*** | 25.5 cm<sup>2</sup> ± 7.5 |
|  | Men | 5.5 cm<sup>2</sup> ± 1.2 | N/A |

### Experiment 1: Tactile pleasantness and intensity via robot-delivered stroking, and thermal thresholds, over body sites

#### Participants

The experiment was conducted on 51 healthy participants (34 females – 17 of which were taking hormonal contraception and 17 were not – and 17 males). Table 1 shows the details of the participants and their TEAQ-s scores. Measure of the areola size are missing for two participants. One participant did not complete the nipple-areola skin site and described being hypersensitive and unable to tolerate it, but did complete the other sites. Due to technical issues, one participant did not provide intensity ratings for the affective touch protocol and another did not provide intensity ratings for the third repetition on the palm for the five velocities nor pleasantness ratings for the third repetition on the palm at velocities 0.3, 3, 10, and 30 cm^-1.^ The missing scores on the third repetition were replaced by the average of the ratings from this participant at the first and second repetitions in each condition.

#### Procedure

Experiments took place in a black room (no window) in which the only openable door was located inside another room to guarantee privacy without key-locking the door. Once the consent forms were signed, participants’ information was collected. Then, the participant was asked to take off their upper-body clothes, put on a custom cotton sheet that had sections that could be undone using resealable openings, and lie down on a massage table (Figure 1). The sheet allowed us to only uncover the skin of the site being tested. When the participant was installed, pillows were used to adjust the position and ensure that the participant was comfortable, and the skin site immobilized. Five skin sites were tested on the left side of the participants, three on the torso: the skin of the upper breast, the skin including the nipple-areola complex, and the skin of the torso below the breast, and two control skin sites: the forearm and the palm. On these skin sites, two experimental protocols were carried out: a classic stroking protocol investigating affective touch and thermal detection thresholds measurements.

**Figure 1:**
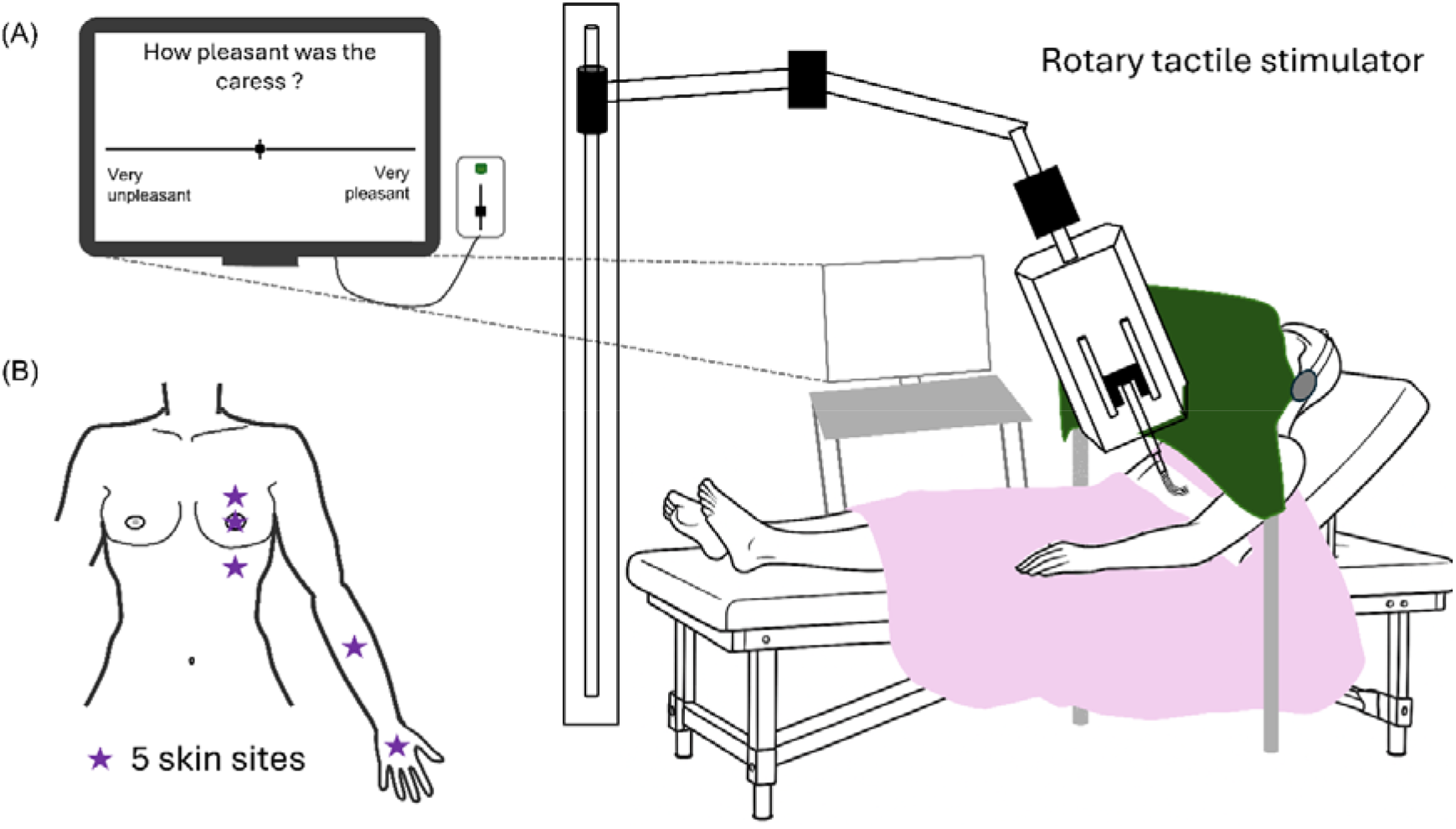
Set-up illustration of the first part of the experiment. (A) Strokes were delivered using a soft brush moved by a rotary tactile stimulator robot and visual analog scales were presented on a computer screen. Participants rated their perception using a slider device. They first rated the pleasantness, from very unpleasant (-10) to very pleasant (+10), then the intensity, from very weak (-10) to very strong (+10). (B) The five skin sites tested, as indicated by stars.

#### Robot-applied stroking touch protocol

A rotary tactile stimulator was used to deliver stroking stimulation with a soft brush at a controlled velocity, direction, and with a calibrated force of 0.4N. The brush was placed approximately 1 cm above the skin, on the center of the skin site, before calibration. A curtain was placed in front of the participant’s face to hide the skin sites and the device from their view, and headphones were placed on the participant’s ears to limit auditory information (Figure 1). Five stroking velocities were tested on each skin site and repeated three times in a pseudo-randomized order; thus, fifteen stimulations were delivered per skin site. The velocities were 0.3 cm. s^-1^ (very slow), 1, 3, 10, and 30 cm.s^-1^ (very fast). The stroking was delivered in a proximal to distal direction on the forearm and palm and in a lateral to medial direction on the torso. Each skin site was tested as a block and sites were tested in a pseudo-randomized order. After each stimulation, the participant used visual analog scales to rate tactile pleasantness, from very unpleasant to very pleasant, and then tactile intensity, from very weak to very strong. The output from the scales ranged from -100 (very unpleasant/very weak) to +100 (very pleasant/very strong) and was converted into -10 to +10 values. The scales were displayed on a monitor on the right of the participants. They used a slider device in their right hand to move a cursor on the screen, and a button to select their response. When ratings were validated, the next stimulation automatically started. There was a pause of at least 30 s between each stroke.

#### Skin temperature and thermal detection thresholds

After testing tactile pleasantness and intensity, the thermal detection thresholds to warm and cool were measured on the same skin sites. A thermode (QST lab) was used with a standard method of limits protocol to apply warmth or cooling to the skin at a temperature rate change of 1°C/s from the baseline temperature of the skin (measured by the instrument, in °C). The participant was instructed to press a button when a temperature change was felt. The warm threshold was measured first, then the cool threshold, both of which were composed of an average of three stimuli, after an initial test stimulus. Thresholds are shown as the difference between baseline temperature and the mean of these three detected temperatures (either warm or cool for the respective thresholds).

### Experiment 2: Tactile pleasantness and intensity via manual-delivered, fully randomized stroking, and tactile sensitivity, over skin sites

In a follow-up experiment, touch perception was investigated using a complementary protocol and tactile sensitivity was investigated by measuring tactile detection thresholds. It was conducted on 18 women and four of these had participated in Experiment 1. The time between participation in each experiment was more than 1 year. Table 1 shows the details of the participants and their TEAQ-s scores. In Experiment 2, a similar stroking protocol was carried out, but all skin sites and stroking velocities were fully randomized. To allow this, the brush stroking was performed by a trained female experimenter (author S.F.), but fewer stroking velocities were used and the stroking was not carried out on the NAC, due to the participant having to rate pleasantness, as delivered by another person, on an erogenous zone, which was considered to cause a potentially uncomfortable situation. Additionally, instead of thermal thresholds as per Experiment 1, tactile detection thresholds were obtained.

#### Experimenter-applied stroking touch protocol

Four of the five previous skin sites were tested: upper breast, torso under the breast, forearm and palm, and three of the previous velocities: 0.3, 3 and 30 cm.s^-1^. Instead of the rotary tactile stimulator robot, a trained experimenter applied the stimulation with an upgraded manual brush and audio-guiding through earphones: an auditory signal was announcing the skin site and the velocity of the next stimulation to apply and playing a rhythm to guide the application of the stimulation. Dots with a pencil were traced on the skin to identify the stimulation area, which was 6 cm long, and were used in combination with the audio rhythm. The force was controlled visually: the experimenter attempted to reproduce the same force applied in each test, causing a similar curvature of the brush hairs. The manual brush was equipped with a gyroscope system (Adafruit BNO055) and with two buttons. The first button was used to move on to the next stimulation and trigger the audio signal (heard by the experimenter only). The second button was pressed by the experimenter during the application of the stimulus, and the angles were recorded throughout this period. Using the recorded angles, the applied velocity was calculated a posteriori for each stimulation. For a target velocity of 0.3 cm.s^-1^, the mean velocity ± 95% CI calculated was 0.66 cm.s^-1^ ± 0.14, for a target velocity of 3 cm.s^-1^, this was 3.26 cm.s^-1^ ± 0.24, and for a target velocity of 30 cm.s^-1^, it was 36.71 cm.s^-1^ ± 3.62. The stimulation order was pseudo-randomized for both the velocity and the skin site. The participant lay down in the same position as in Experiment 1, but could wear a bra. Instead of a curtain and a monitor, the participant wore a virtual reality headset that blocked out the view of the experimenter. The participant saw a screen in a gray environment and after the stimulation was applied, the visual analog scales were displayed: first the pleasantness scale, then the intensity scale, as in Experiment 1.

#### Tactile detection thresholds

After the affective touch protocol, participants could move and relax for a few minutes. Then, the experimenter measured the tactile detection threshold on six skin sites: upper breast, areola, nipple, under the breast, forearm, and palm. Measures were conducted via the application of calibrated monofilaments (Aesthesio set, Ugo Basile). A short familiarization session consisted of applying a first filament (4 g, equivalent to 40 mN) to the center of the participant’s skin site to confirm that they could feel the touch. The participant was asked to indicate each time they felt stimulation (by saying “top”). Monofilaments in the set ranging from 40 mN (4 g) to 0.08 mN (0.008 g) were applied (full range: 20, 14, 10, 6, 4, 1.6, 0.7, 0.4, 0.2, 0.08 mN). An adaptive staircase procedure was used (two-down, one-up), based on the responses made to three applications of a monofilament force, to obtain a representative tactile sensitivity level (Dione et al. 2023; Samain-Aupic et al. 2024). The experiment started with the 40 mN monofilament, except for the palm where it started with 10 mN, as the tactile detection threshold is lower (Ackerley, Carlsson, et al. 2014). If all three applications were felt by the participant, the applied monofilament force level was decreased by two monofilaments (14 mN). If all three applications were felt here, the monofilament force level increased by one (20 mN). This adaptive staircase (two-down, one-up) continued until the participant made an error. If the participant made one error, the monofilament force level was increased by one. The test ended when at least two errors were made at a monofilament level and the force detection threshold was noted as the level just preceding this (i.e. one-up).

### Statistical analysis

For both experiments, analyses were conducted using Python for preprocessing and visualization, and RStudio for statistical analyses. The decimal logarithm transformation of stroking velocity was used to match the methods in the literature (Ackerley, Carlsson, et al. 2014; Löken et al. 2009). A significance threshold of p < 0.05 was used. Ratings of pleasantness and intensity were analyzed using linear mixed-effects models. For each measure, the rating was used as the dependent variable, with the velocity, skin site, and their interaction included as fixed effects. The model included a random intercept and a random slope for the velocity for each subject. Post hoc tests and planned contrasts were performed when appropriate, with p-values adjusted using the Bonferroni correction. In the models applied to the full-group data in Experiment 1, sex and hormonal contraception were also included as fixed effects, with an interaction term for sex. When analyzing data from women in Experiment 1, hormonal contraception was included as a fixed effect with interaction. Baseline temperature was analyzed using a linear mixed-effects model, with the temperature as the dependent variable. Skin site and sex were included as fixed effects with interaction, and hormonal contraception was included as a fixed effect without interaction. A random intercept was included for each participant. Thermal detection thresholds (Experiment 1) and tactile detection thresholds (Experiment 2) were not normally distributed. Thus, non-parametric Friedman (for repeated measures) and Kruskal-Wallis (for group comparisons) tests were used, followed by Wilcoxon signed rank (post Friedman test) and Dunn’s (post Kruskal-Wallis test) post-hoc comparisons with Bonferroni correction.

## Results

### Experiment 1: Tactile pleasantness and intensity via robot-delivered stroking, and thermal thresholds, over body sites

In n = 51 participants, tactile pleasantness and intensity ratings of robot-applied brush stroking were analyzed separately using regression analyses of the ratings over the stroking velocities at each skin site, testing linear and quadratic fit models, and linear mixed models with follow-up Bonferroni-corrected tests.

#### Tactile pleasantness ratings

The pleasantness ratings over the stroking velocities from 0.3 to 30 cm.s^-1^ followed inverted U-shaped relationships for all skin sites, where pleasantness peaked around the intermediate velocities. A linear mixed model on all participants revealed a significant effect of stroking velocity (F_4,63.41_ = 41.39, p < 0.001, η^2^ ⍰ = 0.72 ), skin site (F_4,1024.03_ = 2.80, p = 0.025, η^2^⍰ = 0.01 ), and no effect of the participant sex (p = 0.085), or hormonal contraception (p=0.097), but a significant interaction between skin site and sex (F_4,1024.03_ = 6.92, p < 0.001, η^2^⍰ = 0.03). Due to the interaction between skin site and sex, further analyses were conducted on the data from women and men separately.

For women (n = 34), all mean pleasantness curves per skin site exhibited significant negative quadratic regressions (all p < 0.001), and no significant linear fits were found. In a linear mixed model, we found a significant effect of velocity (F_4, 41.40_ = 36.70, p < 0.001, η^2^⍰ = 0.78), skin site (F_4, 667.33_ = 3.77, p = 0.005, η^2^⍰ = 0.02), and an interaction between the velocity and the skin site (F_16, 667.72_ = 2.26, p = 0.003, η^2^⍰ = 0.05), but no effect of hormonal contraception (p = 0.073), nor the interactions between stroking velocity and hormonal contraception (p = 0.505), skin site and hormonal contraception (p = 0.242), nor between the three (p = 0.940) (Figure 2A). Post-hoc tests comparing different stroking velocities found differences between 0.3 compared to 1, 3, and 10 cm.s^-1^ (all p < 0.001), and between 30 compared to 1, 3, and 10 cm.s^-1^ (all p < 0.001). Post-hoc tests comparing skin sites found that the NAC was significantly less pleasant than the forearm (p = 0.009) and the palm (p = 0.001). Follow-up contrast comparisons between the skin sites at each velocity revealed significant differences specifically at 30 cm.s^-1^, where the NAC was rated less pleasant than the palm (p < 0.001), forearm (p < 0.001), and upper breast (p = 0.039). Further, stroking under the breast was rated as less pleasant than the forearm (p = 0.013).

**Figure 2:**
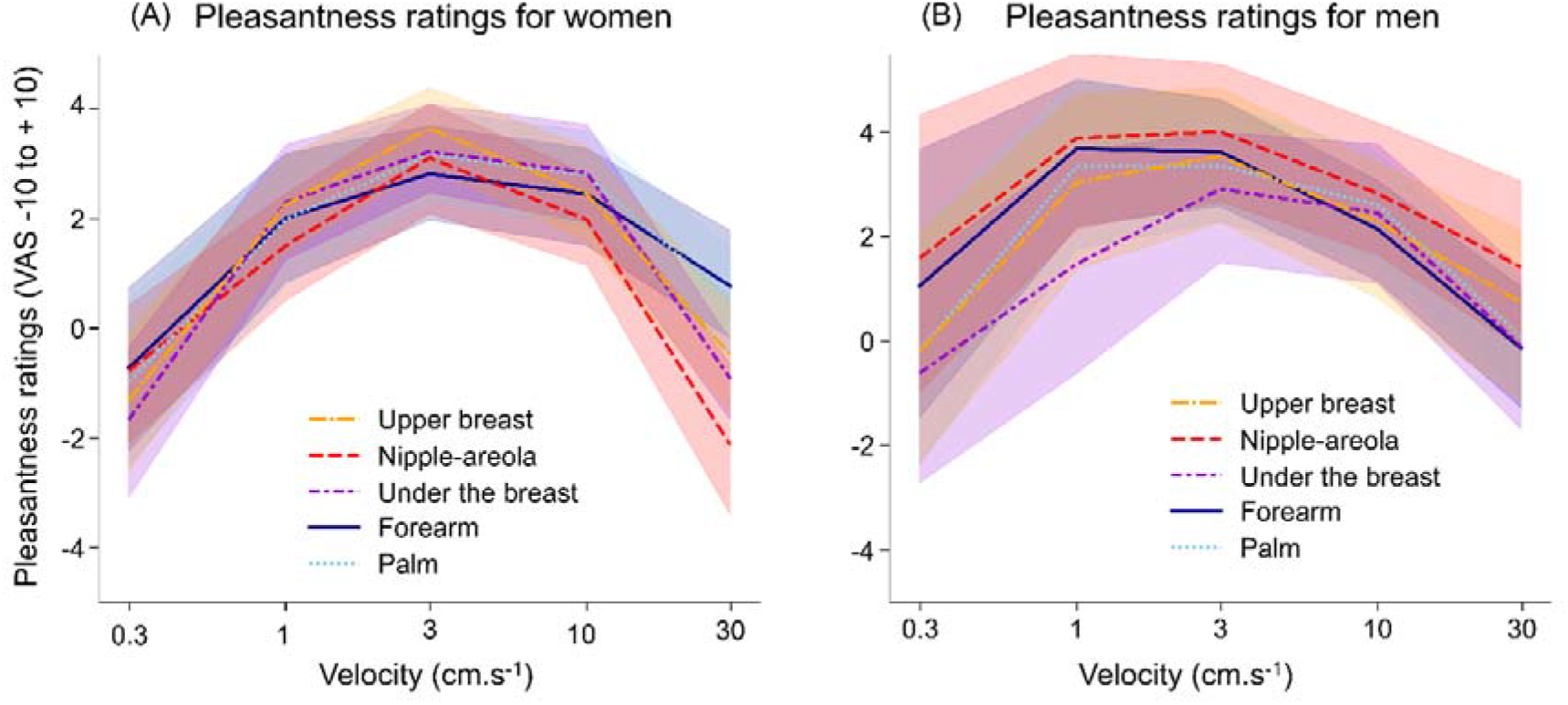
Pleasantness ratings over stroking velocity, per skin site (Experiment 1, robot-applied). Mean pleasantness ratings over stroking velocities for each skin site from robot-applied stroking in Experiment 1 for (A) women (n = 34) and (B) men (n = 17). The full rating visual analog scale (VAS) was from -10 (very unpleasant) to +10 (very pleasant). Error bands correspond to 95% confidence intervals.

For men (n = 17), all mean pleasantness curves exhibited significant negative quadratic regressions (p = 0.037 for the NAC, p < 0.01 for the other skin sites), and no significant linear fits were found. A linear mixed model found significant main effects of stroking velocity (F_4,27.18_ = 14.88, p < 0.001, η^2^⍰ = 0.69) and skin site (F_4,352.02_ = 5.24, p < 0.001, η^2^⍰ = 0.06), but no significant interaction (p = 0.602) (Figure 2B). Post-hoc tests between velocities found differences between 0.3 compared to 1 and 3 cm.s^-1^ (p = 0.003 and p = 0.040, respectively) and between 30 compared to 1 (p = 0.040), 3 and 10 cm.s^-1^ (both p < 0.001). Post-hoc tests between pleasantness ratings across skin sites showed that under the breast was significantly rated less pleasant than the NAC (p<0.001).

#### Tactile intensity ratings

The tactile intensity ratings over stroking velocities showed a clear positive linear relationship, where increased speed increased the sensation. All intensity relationships per skin site exhibited significant linear regressions (all p < 0.001). A linear mixed model on the results from all participants revealed a significant effect of stroking velocity (F_4, 83.73_ = 5.22, p < 0.001, η^2^⍰ = 0.20) and skin site (F_4, 1006.79_ = 7.51, p < 0.001, η^2^⍰ = 0.03), but no effects of the sex (p = 0.150), hormonal contraception (p = 0.153), or the interactions between the velocity and skin site (p = 0.226), the velocity and the sex (p = 0.945), the skin site and the sex (p = 0.151) or between the three (p = 0.305) (Figure 3). Bonferroni corrected post-hoc comparisons showed that all the velocities were significant (all p < 0.05, most p < 0.01), and that the NAC was more intense than the forearm (p = 0.010) and under the breast (p = 0.016), while the palm was more intense than the forearm and under the breast ( both p <0.001).

**Figure 3:**
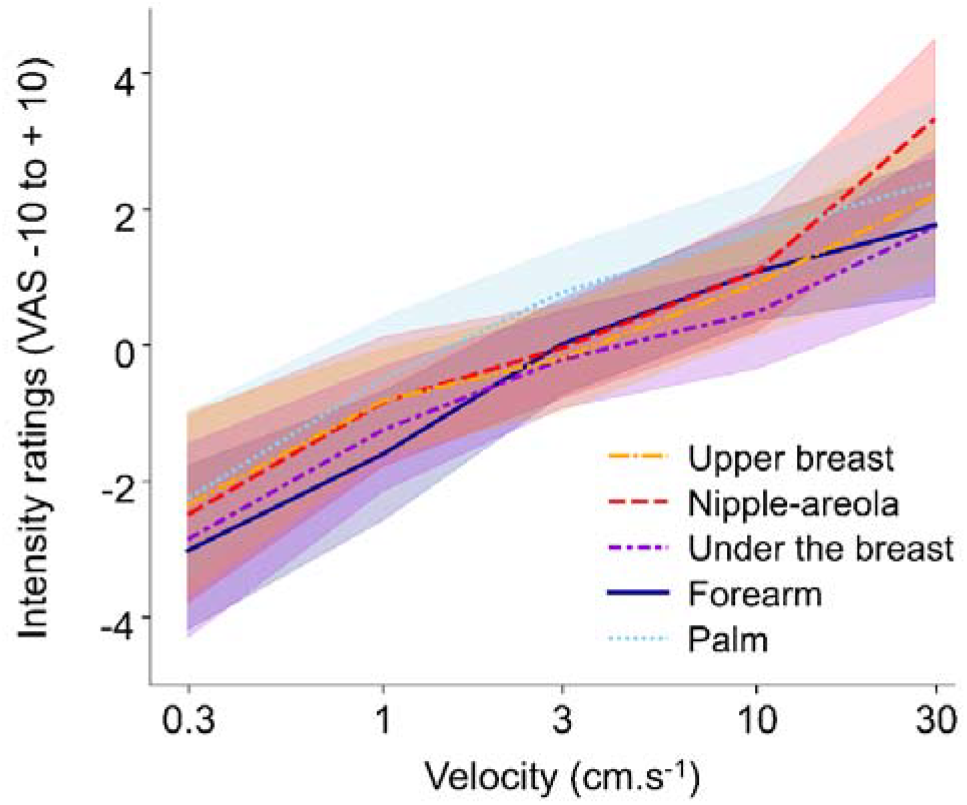
Intensity ratings over increasing stroking velocity, per skin site (Experiment 1, robot-applied). Mean intensity ratings over stroking velocities for each skin site from robot-applied stroking in Experiment 1. The full rating visual analog scale (VAS) was from - 10 (very weak) to +10 (very strong) (n = 51 participants). Error bands correspond to 95% confidence intervals.

#### Skin temperature and thermal perception

The baseline temperature measured across the skin sites was determined before temperature threshold detection testing (Figure 4A). A linear mixed model revealed an effect of the skin site (F_4, 196_ = 14.70, p < 0.001, η^2^⍰ = 0.23) and an interaction between the skin site and the sex (F_4, 196_ = 6.41, p < 0.001, η^2^⍰ = 0.12), but no effect of the sex (p = 0.463) or hormonal contraception (p = 0.683). Bonferroni-corrected post hoc comparison found the NAC baseline temperature to be higher than the palm and the forearm (both p < 0.001), and the upper breast to be higher than the palm and the forearm (both p <0.001) and under the breast (p = 0.009). Post-hoc comparison on the sex found no significant difference, but further contrast comparisons found a significant difference on the palm between men and women (p = 0.004).

**Figure 4:**
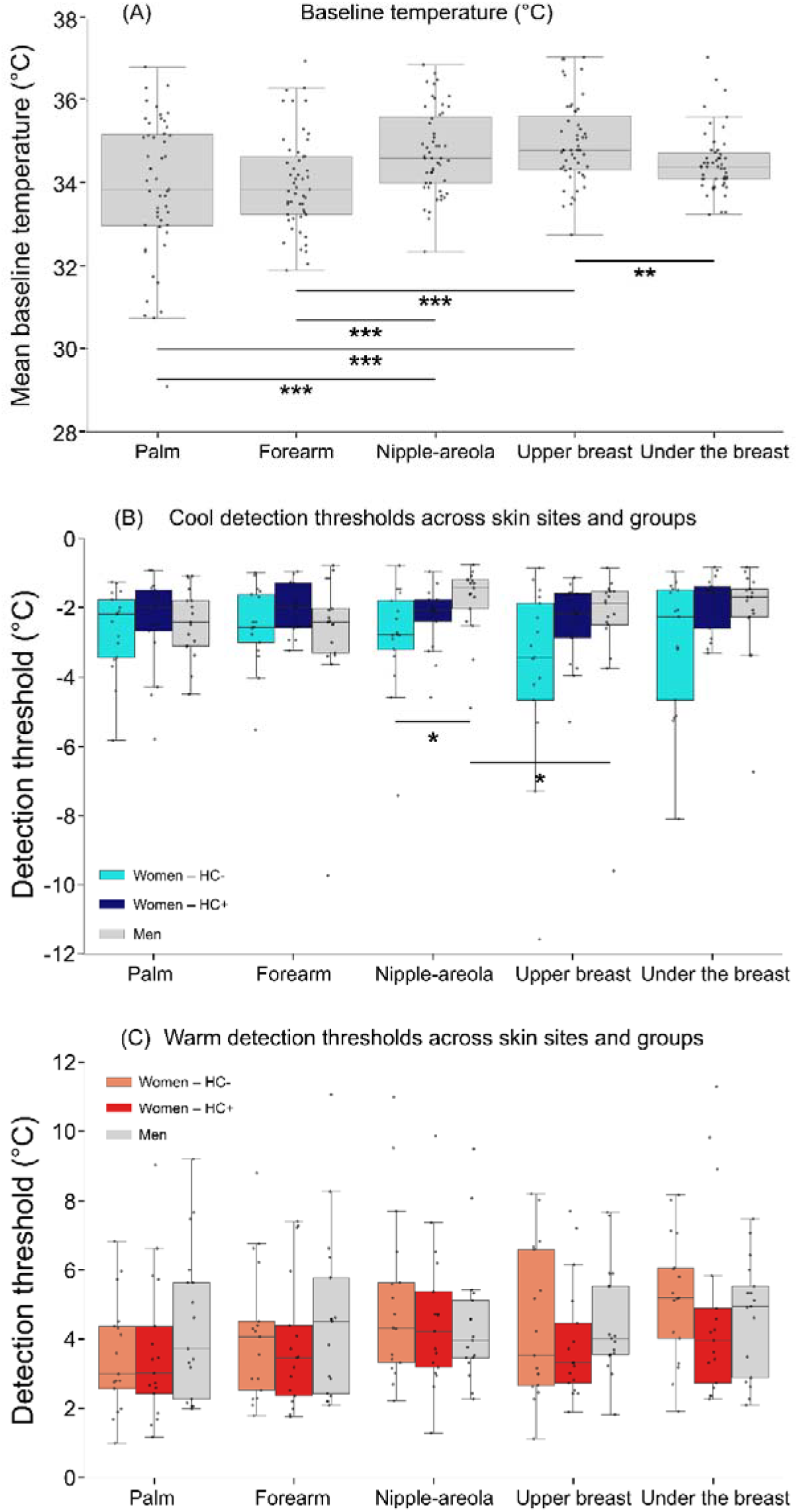
Baseline skin temperature and cool and warm detection thresholds over skin sites (Experiment 1). Boxplots of (A) mean baseline skin temperatures, (B) cool detection thresholds, and (C) warm detection thresholds per skin site. For the cool and warm detection thresholds, participants are presented as three groups (n = 17 per group). ‘Women – HC+’ corresponds to the group of women who are currently taking hormonal contraception, ‘Women – HC-’ corresponds to the group of women who are not currently taking hormonal contraception, and ‘Men’ to the male participants. Significant effects: ^**^p < 0.01, ^***^p < 0.001.

We then investigated thermal detection thresholds between skin sites and participant groups. Thresholds were calculated as the difference between the baseline skin temperature and the mean of the temperature eliciting warm or cool perception from three tests each. As the data were not normally distributed, we used non-parametric tests and also separated the comparisons by group, per skin site, due to the interaction that was found in the baseline skin temperature. Groups were differentiated in men (n = 17), women who are currently taking hormonal contraception (HC+, n = 17), and women who are not currently taking hormonal contraception (HC-, n = 17).

For the cool detection thresholds, significant effects of skin site (χ^2^(4) = 15.823, p = 0.003) and group (χ^2^(2) = 12.207, p = 0.002) were found (Figure 4B). Bonferroni-corrected post-hoc tests for skin site differences showed that the NAC had a smaller cool detection threshold than the upper breast (p = 0.037). In terms of group differences, women who were not taking hormonal contraception had significantly larger detection thresholds than both women taking hormonal contraception (p = 0.009) and men (p = 0.006). Further, a lower cool detection threshold was found at the NAC skin site in men compared to the NAC in women who were not using hormonal contraception (p = 0.015) and compared to the upper breast skin site in men (p = 0.028). For warm detection thresholds (Figure 4C), there was a significant effect of skin site (χ(4) = 12.67, p = 0.013), but no effect of group (p = 0.224); however, no significant post-hoc comparisons were found using Bonferroni-corrected post-hoc tests.

#### Correlations between TEAQ-s scores and the other measures

We collected scores from the TEAQ-s (Touch Experience and Attitudes Questionnaire – short version) to look at correlations with all the measures. FDR-corrected Spearman correlations showed a significant positive correlation between pleasantness ratings and TEAQ-s scores (ρ = 0.40, p = 0.019), i.e. more positive experiences and attitudes toward touch were correlated with higher pleasantness ratings. We also found a positive correlation between absolute warm and cool detection thresholds (ρ = 0.52, p = 0.001), which can be interpreted as higher warm detection thresholds being correlated with lower cool detection thresholds, i.e. the less sensitive participants were to warm detection, the less sensitive they were to cold detection. In addition, we also found a negative correlation between absolute cool detection thresholds and intensity ratings (ρ = -0.39, p = 0.019), meaning that better cool detection (less deviation from skin temperature) was related to higher sensations of tactile intensity.

### Experiment 2: Tactile pleasantness and intensity via manual-delivered, fully randomized stroking, and tactile sensitivity, over skin sites

We further investigated the effect of skin site, due to the unavoidable experimental confound that the stroking was delivered to the skin sites in blocks using the robot in Experiment 1. Thus, Experiment 2 used manual stroking of the skin at the velocities of 0.3, 3, and 30 cm.s^-1^ in n = 18 women and was fully randomized between stroking velocity and skin site. Due to the more sensitive nature of experimenter-delivered brush strokes, the participants were not stroked on the NAC. Additionally, instead of obtaining thermal perception thresholds, we obtained tactile detection thresholds over all skin sites, including the nipple and areola separately.

#### Tactile pleasantness ratings

In Experiment 2, with the fully randomized protocol, mean pleasantness rating over the stroking velocity still followed an inverted U-shaped curve for all the skin sites. All curves exhibited significant negative quadratic regressions per body site (all p < 0.001) and no significant linear regressions were found (all p > 0.2) (Fig. 5A). A linear mixed model on the results revealed a significant effect of the velocity (F_2,17.22_ = 22.25, p < 0.001, η^2^⍰ = 0.72; Fig. 5B) and of the skin site (F_3,153.78_ = 4.99, p < 0.01, η^2^⍰ = 0.09; Fig. 5C), but no significant interaction (p = 0.577). Bonferroni-corrected post-hoc comparisons showed that 3 cm.s^-1^ was rated more pleasant than 0.3 cm.s^-1^ and 30 cm.s^-1^ (both p < 0.001). For the skin sites, post-hoc tests revealed that under the breast was significantly less pleasant than the forearm (p = 0.026) and the upper breast (p = 0.020).

**Figure 5:**
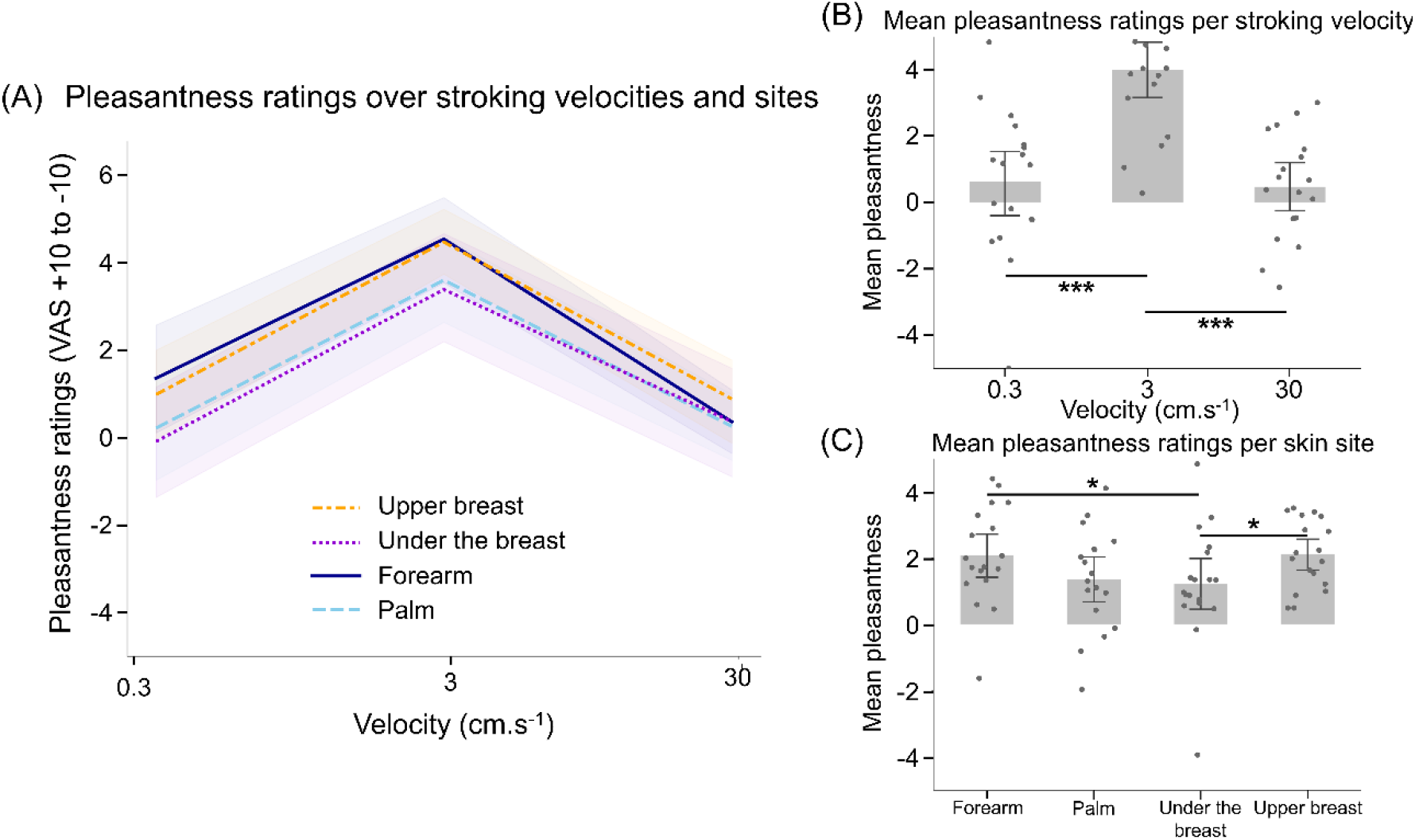
Pleasantness ratings over increasing stroking velocities, per skin site (Experiment 2, experimenter-applied). (A) Pleasantness ratings over stroking velocities for each skin site, (B) mean ratings for each skin site and (C) for each stroking velocity (n = 18 women). The full rating visual analog scale (VAS) was from -10 (very unpleasant) to +10 (very pleasant). Error bands and bars correspond to a 95% confidence interval. Significant effects: ^***^p < 0.001.

#### Tactile intensity ratings

For the experimenter-applied intensity ratings, all regressions exhibited significant linear fits (all p < 0.001) (Fig. 6A). A linear mixed model showed a significant effect of stroking velocity (F_2,17_ = 13.33, p < 0.001, η^2^⍰ = 0.61; Fig. 6B) and skin site (F_3,153_ = 15.90, p < 0.001, η^2^⍰ = 0.24; Fig. 6C), but no significant interaction (p = 0.358). Using post-hoc comparisons by velocity, 30 cm.s^-1^ was rated as more intense than 0.3 cm.s^-1^ (p < 0.001) and 3 cm.s^-1^ (p = 0.002), and 3 cm.s^-1^ was rated more intense than 0.3 cm.s^-1^ (p = 0.002). On the skin site, post-hoc comparisons revealed that the palm was perceived as more intense than the forearm, the upper breast (both p < 0.001), and under the breast (p = 0.001).

**Figure 6:**
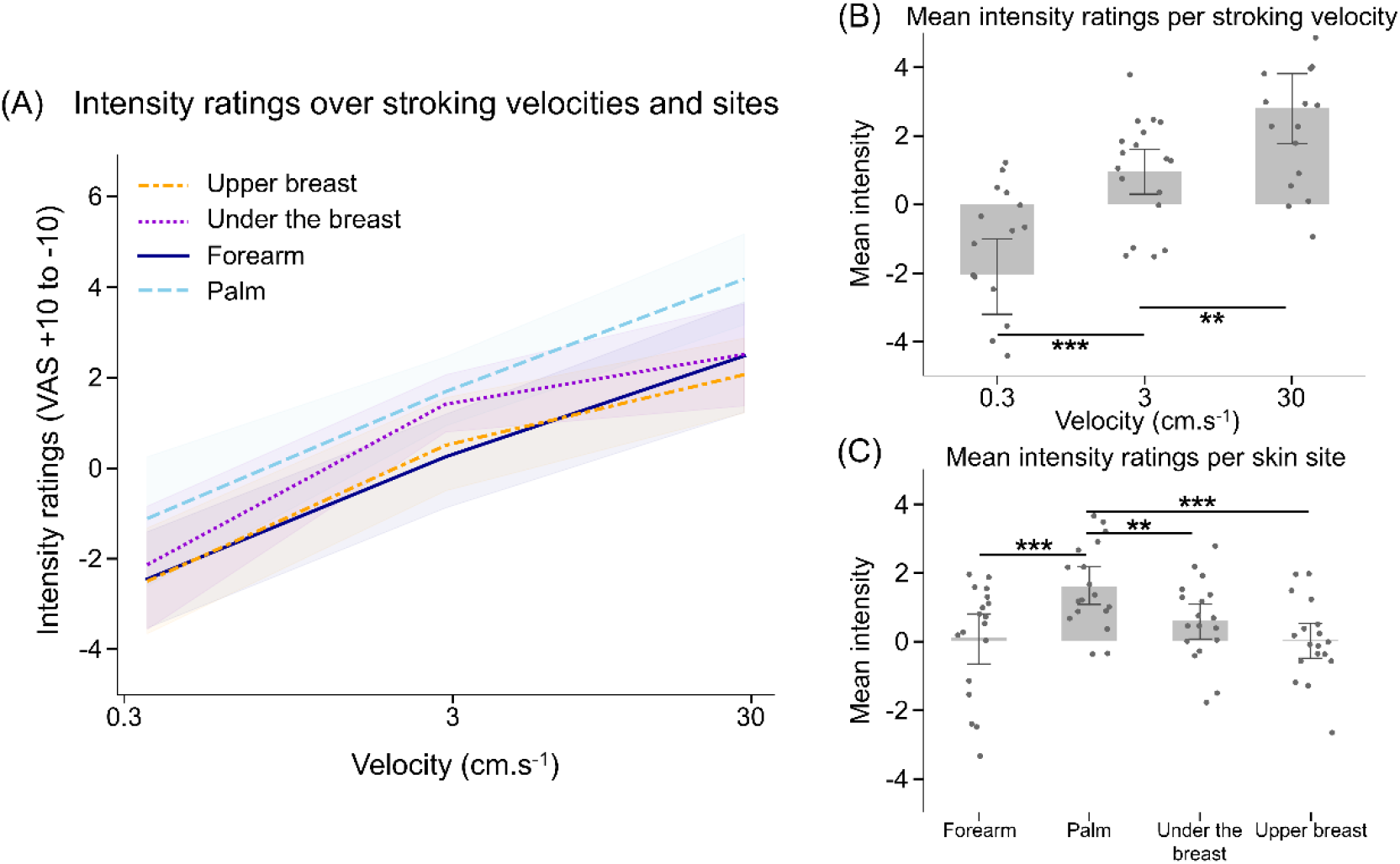
Intensity ratings over increasing stroking velocities, per skin site (Experiment 2, experimenter-applied). (A) Intensity ratings over stroking velocities for each skin site, (B) mean ratings for each skin site and (C) for each stroking velocity (n = 18 women). The full rating visual analog scale (VAS) was from -10 (very weak) to +10 (very strong). Error bands and bars represent 95% confidence intervals. Significant effects: ^*^p < 0.05, ^**^p<0.01, ^***^p < 0.001.

#### Tactile sensitivity

To complement the previous experiment, in Experiment 2, we obtained measures of tactile sensitivity, as measured by the detection threshold from calibrated monofilaments. We found that this differed across skin sites (χ^2^(5)= 48.91, p < 0.001) (Fig. 7). Bonferroni-corrected post-hoc comparisons revealed that the areola was less sensitive than all the other sites: forearm (p = 0.007), nipple (p = 0.019), palm (p = 0.003), under the breast (p = 0.005), and upper breast (p = 0.010). The upper breast was also less sensitive than the palm (p = 0.013).

**Figure 7:**
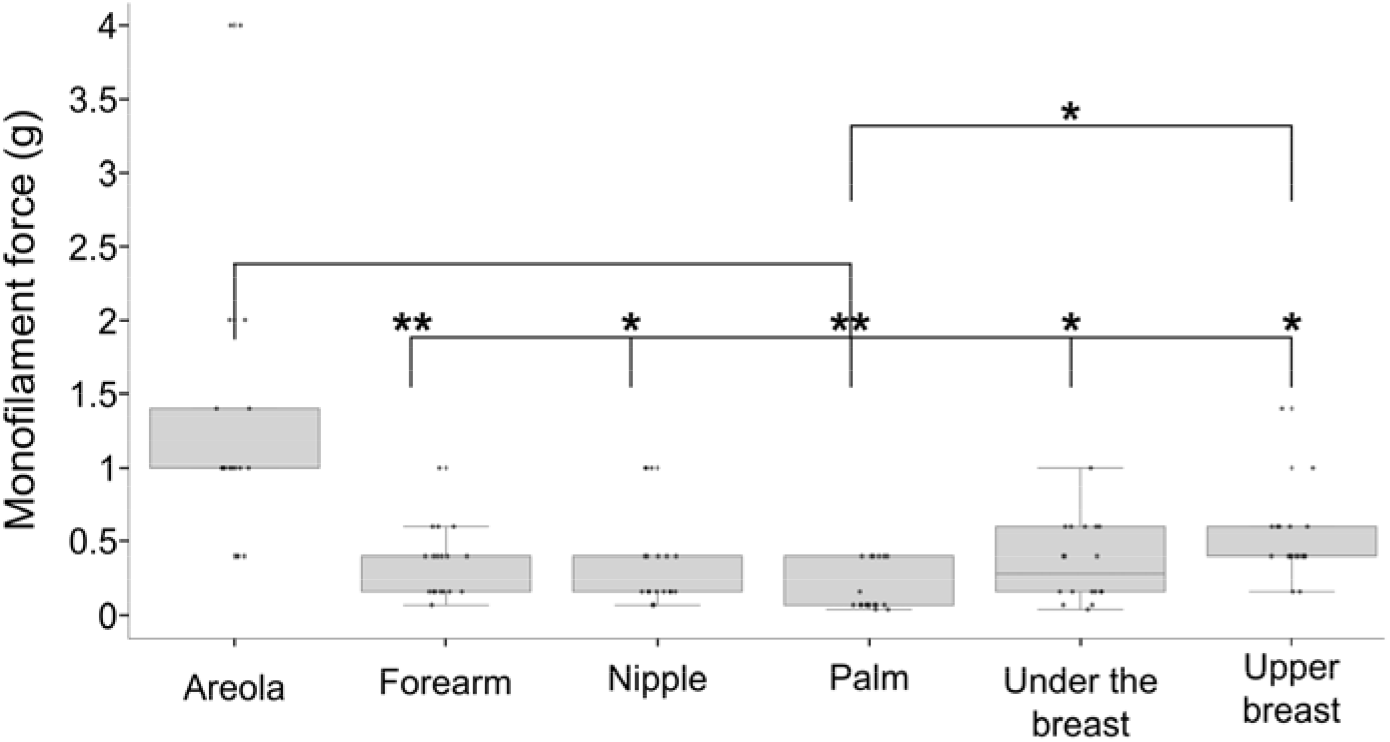
Tactile sensitivity over skin sites, measured by calibrated monofilament detection (Experiment 2). Boxplots of the tactile detection thresholds measured over skin sites using a range of forces applied with calibrated monofilament indentation (n = 18 women). Significant effects were found: ^*^p < 0.05, ^**^p < 0.01.

## Discussion

Our present results, over both experiments, highlight that the torso shares similarities in tactile perception with the more comprehensively studied skin sites of the forearm and palm, but also has specific perceptual characteristics. The perception of touch on the torso seems to somewhat differ between the breasts and the rest of the torso, and across the breast, where tactile sensations varied between the hairy skin of the upper breast, the mixed skin of the areola, and the hairless skin of the nipple, suggesting that these are three distinct skin areas that give rise to different percepts.

Concerning pleasant touch perception, we found that this followed the classic inverted U-shaped curve over increasing stroking velocity on the breast and torso, as well as on the forearm and palm. This relates well to the literature showing that the inverted-U-shaped pleasantness curve seems to be relatively similar and stable over face and limb skin sites (Ackerley, Carlsson, et al. 2014; Cazzato et al. 2021; Jönsson et al. 2015; Cruciani et al. 2021). Our additional sites on the torso, namely the upper breast, NAC, and under the breast, did not show any major increases or decreases in pleasantness, but we did find subtle differences on the NAC in women and on the skin under the breast for all participants. Specifically, the difference was more profound at the fastest stroking velocity, which is often where distinctions between conditions are found (Ackerley, Carlsson, et al. 2014; Faresse et al. 2026). The slightly less pleasant perception under the breast is consistent with two previous studies that showed that the abdomen was perceived as less pleasant than the other body sites tested in similar stroking touch protocols (compared to the palm, forearm, face, and back (Cazzato et al. 2021), and compared to the forearm and back (Wyder-Hodge 2025)). In addition to these similarities, in the first experiment, the NAC was perceived as less pleasant, especially at the highest stroking velocity, in women, but not in men, where it was more pleasant than under the breast. Breasts are erogenous zones (Nummenmaa et al. 2016), with morphological differences between men and females, and the nipples are of particular importance in sexual function and arousal, especially in women (Levin and Meston 2006; Lindau et al. 2023). On the contrary, erogenous touch on the penis appears to be different, as a study found that the stroking touch stimulation delivered by a partner on the penis was overall more pleasant than the forearm (Ruesink et al. 2022). Whether these results are related to C-LTMR or other mechanoreceptor innervation remains to be determined.

The inverted-U-shape pleasantness curve has been correlated in literature with the activity of C-LTMRs (Löken et al. 2009; Ackerley, Wasling, et al. 2014), and our results reproduced this curve on all skin sites. This correlation between C-LTMR firing and pleasantness has only been established on the hairy skin of the forearm, therefore it is not known whether this relates to the density or mere presence of C-LTMRs over other skin sites. In general, pleasantness of stroking touch on the leg has been found to be very similar to that on the forearm (Ackerley, Carlsson, et al. 2014; Jönsson et al. 2015), although the density of C-LTMRs in the human leg is rather low (Löken et al. 2022). Similarly, pleasant stroking on the face has been found to be similar (Ackerley, Carlsson, et al. 2014) or even more pleasant (Essick et al. 1999) than on the forearm, even though there is a lower density of C-LTMRs on the face than on the forearm (Nordin 1990; Vallbo et al. 1999). Further, C-LTMRs are found on the glabrous hand skin, yet only rarely (Watkins et al. 2021), but the corresponding pleasantness curve on the palm is typically similar to the forearm (Ackerley, Carlsson, et al. 2014; Cruciani et al. 2021), as we found presently. Therefore, the evidence suggests that C-LTMR presence may be sufficient to influence the inverted-U shape pleasantness curve, but that this curve may not necessarily be related to the density of C-LTMRs. Even so, evidence in the literature supports the contribution of other mechanoreceptors in affective touch (Vallbo et al. 2009; Schirmer et al. 2023; Case et al. 2023).

We found a linear curve for tactile intensity over increasing stroking velocity, which relates more to the activity of all types of Aβ mechanoreceptors. We found that non-hairy skin sites (palm and NAC) were consistently perceived as more intense. This may suggest a higher density of mechanoreceptors, differences in the types of mechanoreceptors present (e.g. Krause corpuscles), or a difference in the balance of mechanoreceptors. Tactile intensity has been associated with the activation of Aβ mechanoreceptors (Löken et al. 2009; Edin et al. 1995), which are dense in the glabrous skin of the hand (Corniani and Saal 2020). However, it has been shown that the skin of the nipple is not densely innervated with Aβ mechanoreceptors (Gutiérrez-Villanueva et al. 2020), suggesting that the increase in intensity perceived at the NAC at 30 cm.s^-1^ may have a different origin, for example, the potential presence of other types of mechanoreceptor (e.g. Krause corpuscles that are found in mucocutaneous skin (Winkelmann 1957)). As well as the results using moving touch, we also found that tactile sensitivity on the areola was significantly worse than on all the other skin sites. The tactile sensitivity tests were only carried out in Experiment 2, thus only in women, but it demonstrates a poor tactile sensitivity of areola, while the nipple was very sensitive. This suggests that even if often considered as a complex, the NAC may include two different skin types: the nipple and the areola, which have different innervations by sensory afferents.

Concerning the sex differences that we found, namely on the NAC for pleasantness and cool perception, these could be due to differences in the sizes of the NAC between sexes and/or the effect of hormones. Our measures of NAC area between men and women in Experiment 1 showed that women had significantly larger NACs than men, which could mean differences in sensitivity. The NAC area is likely larger in women due to hormonal influences, such as increased breast size and the NAC growing in-line with the breast. However, a larger NAC would potentially mean less sensitivity, as seen with the cool detection thresholds, due to a lower density of receptors over the larger area, similar to the effect seen on the fingertip for tactile spatial discrimination (Peters et al. 2009). A previous study in women also found differences in cool sensitivity across the NAC, but not for warmth (Blount et al. 2025). As we only tested women in Experiment 2, we cannot say if there are sex differences in tactile detection on the nipple or areola, but a previous study found that females were generally more sensitive to touch, including on the breast (Weinstein 1968), which does not align with larger breasts being less sensitive to touch. Our finding that stroking touch over the NAC was less pleasant for women may relate to the sensitivity of the NAC and the combination of hormonal influences.

As well as general hormonal differences, there could also be differences due to the effect of the menstrual cycle. A study has shown that the detection of high-frequency vibrations varied during women’s menstrual cycle, with increased sensitivity during the premenstrual period compared to the postmenstrual period (Gescheider et al. 1984). Further, the study showed that women taking hormonal contraception had stable and highly sensitive thresholds throughout their cycle, similar to those of the premenstrual threshold in women not on hormonal contraception, whereas men had stable thresholds, but these were similar to the less-sensitive post-menstrual thresholds of women not taking hormonal contraception. Due to the specific effects only found at high frequency tactile vibrations, the increased sensitivity was attributed to the influence of female hormones on the Pacinian system, peripherally and/or centrally. Therefore, it is difficult to speculate the extent to which the menstrual cycle effects other aspects of touch, given the specificity of this finding and that we did not currently control for hormonal cycle, yet this is of interest to pursue.

Our measures of the baseline skin temperature showed that the skin sites on the breast (upper breast and nipple areola) were warmer than the other sites. This is consistent with a previous study that investigated thermal perception on the whole body and found that the chest has higher skin temperature (Luo et al. 2020). In another study, thermal perception on the breast (excluding the NAC) and pelvis was investigated and the temperature of the skin under the breast was found to be the highest of the skin sites tested (Valenza et al. 2023), whereas in our study, it was lower than that of the upper breast. During our experiment, the torso skin sites were covered by a sheet between the tests, to respect the intimacy of the participants. This may have influenced the difference observed with the skin temperature of the hand and forearm, which were not covered, but it does not explain the difference observed between the upper part of the breast and the skin under the breast. Furthermore, wearing clothing on this part of the body is typical, suggesting that our results may, to some extent, be similar to those that could be observed under everyday physiological conditions. We also found a difference in baseline temperature between men and women on the palm, where women had lower baseline temperature than men, which has been previously linked to lower heat production in women, due to a generally smaller body size (Yanovich et al. 2020).

Thermoregulation has been found to be influenced by the hormonal cycle in women, being higher post-ovulation than pre-ovulation (Baker et al. 2020). In addition, we found sex differences in cool, but not warm, detection thresholds at the NAC (Blount et al. 2025), which was more sensitive in men than in women who were not taking hormonal contraception. Several factors, as outlined above, may influence the perception, but the difference with women who were not taking hormonal contraception suggests a hormonal influence. In support of this, previous work has shown that cold detection thresholds have been found to differ over skin sites during the menstrual cycle (Söderberg et al. 2006). Other results that we found included a significative positive correlation between absolute cool and warm detection thresholds, indicating that participants who had more sensitive cool detection thresholds also had more sensitive warm detection thresholds (Valenza et al. 2023). We also found a correlation between cool detection thresholds and intensity ratings, where better cool sensitivity was linked to better tactile sensitivity. In addition to the sensory measures, we found a correlation between average pleasantness ratings and the overall TEAQ-s score in Experiment 1, suggesting a link between positive experiences and attitudes toward touch and the overall perception of the pleasantness of stroking touch. This relates well to work showing that increased exposure to inter-personal touch is related to more sensitive pleasant touch appreciation (Sailer and Ackerley 2019). Further, in a study examining the distribution of acceptability of touch across the body according to emotional bonds, found a relation between the strength of emotional bond and ‘touchability’ of the torso (Suvilehto et al. 2015). Overall, these are in line with the affective touch hypothesis where it is proposed that the perception of affective touch is complex and mediated by both peripheral and central mechanisms (Vallbo et al. 2009; Schirmer et al. 2023).

In our present study, we tested sensitivity over two experiments, to gain further insights into sensory processing. A previous study directly comparing stroking delivered by a robot or by an experimenter showed similar pleasantness perception on the forearm, although experimenter-applied stroking was more intense than robot-applied, which was potentially attributed to differences in the application (Triscoli et al. 2013). Although we did not compare these directly in the same experiment, the VAS ratings were very similar for pleasantness and intensity between experiments. Further, we conducted Experiment 2 as a control that randomized skin site, which we were not able to do in Experiment 1, as well as to gain additional tactile detection thresholds. We found no evidence for an effect of blocking the stroking for skin site, although previous work did show some effect between the palm and forearm (Löken et al. 2011), thus further work could investigate this specifically.

Overall, more research is needed to better understand the commonalities and specificities of innervation in the breast and torso. Recording the activity of afferent fibers from the intercostal nerves, which are primarily involved in the innervation of the torso (Chin et al. 2021), or other nerves that contribute to it, such as the supraclavicular nerves, would deepen our understanding by precisely identifying the afferent fibers present and their properties. Given the technical challenge this would represent to conduct microneurography on such small nerves on the torso, where breathing movements would likely affect the recordings, complementary psychophysical approaches would allow for further investigation of the factors that may influence tactile perception in this region of the body. Further, there remain many questions about how the tactile and thermal sensitivity of the breast is influenced by hormones in general and over the menstrual cycle. In conclusion, we find that tactile perception specificities on the torso deserve careful consideration, particularly considering the emerging field of neuroprosthetics, where efforts are being made to improve breast sensation in people who have undergone mastectomy, a procedure that remains relatively common in the treatment of breast cancer, affecting millions of women worldwide (Lindau and Bensmaia 2020). Our results therefore help shape neuroprosthetic innovation for breast peripheral nerve implants by showing that females have areas of the breast that are more sensitive that others, that touch can be intense and thus become unpleasant, and the possibility for hormonal influences on sensory perception.

## Acknowledgements

S.F. was funded by the CNRS MITI program in a grant to R.A. We thank Sliman Bensmaia for his scientific input in conceiving the experiments.

## Authors contributions

S.F., R.H. and R.A. designed the experiment and S.F. and R.A. performed, and analyzed the data. S.F. drafted the first version of the manuscript. All authors approved the final version of the manuscript. All authors agree to be accountable for all aspects of the work in ensuring that questions related to the accuracy or integrity of any part of the work are appropriately investigated and resolved. All persons designated as authors qualify for authorship, and all those who qualify for authorship are listed.

## Data availability

Data and scripts are available via an OSF repository at: https://osf.io/rkd74/overview?view_only=2ff2bd783edc4116ae16fb94114ced90

## Additional information

Competing interests: All authors report no conflict of interest.

